# Loss of adenylosuccinate synthetase 1 in mice recapitulates features of *ADSS1* myopathy

**DOI:** 10.1101/2025.10.01.679796

**Authors:** Morgan E. Kim, Kathryn M. Yammine, Emily T. Hickey, Catalina Matias, Lou C. Dubosclard, Jeffrey J. Widrick, Jeffrey J. Brault, Behzad Moghadaszadeh, Alan H. Beggs

**Affiliations:** The Manton Center for Orphan Disease Research, Division of Genetics and Genomics, Boston Children’s Hospital, Harvard Medical School, Boston, MA, USA; Faculty of Science & Engineering, Sorbonne Université, Paris, France; Indiana Center for Musculoskeletal Health, Department of Anatomy, Cell Biology & Physiology, Indiana University School of Medicine, Indianapolis, IN, USA

## Abstract

*ADSS1* myopathy is an ultra-rare congenital myopathy characterized by progressive cardiac and skeletal muscle degeneration with childhood to adolescent onset. This autosomal recessive disease is caused by mutations in the *ADSS1* gene, encoding the enzyme adenylosuccinate synthetase (AdSS1). AdSS1 plays a critical role in the adenine nucleotide cycle, which is important for energy metabolism in muscle cells. Enzymatic defects, engendered by loss-of-function mutations in *ADSS1*, lead to a bottleneck in the adenine nucleotide cycle, causing metabolic dysfunction which ultimately results in progressive muscle weakness, mobility impairment, and respiratory and cardiac dysfunction, often requiring the use of a ventilator. Despite its debilitating nature, there are currently no cures or targeted treatments available, and little research into possible therapeutic strategies has been done. With a limited patient profile encompassing fewer than 200 known patients worldwide, establishing a mouse model for *ADSS1* myopathy is critical to understanding its pathogenesis and for developing future therapies. Here, we present and characterize the first mouse model of *ADSS1* myopathy – a constitutive *Adss1* knockout model – by (1) defining its natural history, (2) exploring its metabolic pathomechanisms, and (3) characterizing its histopathological features. We find that *Adss1*^KO/KO^ mice have subtle motor deficits and present with histopathological features consistent with patient phenotypes. Overall, we show that despite a relatively mild phenotype, this novel mouse model has quantifiable pathological features that can be used to develop therapies for, and further probe pathophysiology of, *ADSS1* myopathy.

## INTRODUCTION

*ADSS1* myopathy (OMIM 617030) is a recently reported ultrarare myopathy caused by mutations in the *ADSS1* gene, encoding the enzyme adenylosuccinate synthetase 1 (AdSS1).^1^ Genetically, it is an autosomal recessive disease caused by various loss-of-function mutations spanning the coding region.^1^ First reported in Korea in 2016,^1^ the majority of patients identified are of East or South Asian descent, with the most common disease-causing variant arising from a common founder mutation.^2^ Clinically, patients exhibit proximal and/or distal weakness in lower limb muscles with generalized muscle atrophy, fatigue, dysphagia and some masticatory dysfunction. Severe cases may also include cardiomyopathy or diaphragmatic atrophy with reliance on ventilatory support.^2, 3^ Currently, there are no treatments or targeted therapies.

Biochemically, AdSS isoforms play a major role in metabolism by converting inosine monophosphate (IMP) into adenosine monophosphate (AMP) via adenylosuccinate (s-AMP).^4^ The enzyme is found in all life forms with few exceptions.^4^ Two isoforms exist in vertebrates, AdSS1 which is highly expressed in striated muscles, and AdSS2, which is broadly expressed across many tissues at low levels.^5^ AdSS1 serves a critical role in muscle energetics by catalyzing the first and rate limiting step in the conversion of IMP into AMP within the purine nucleotide cycle.^6^ This cycle is linked to the purine salvage pathway and *de novo* purine biosynthesis, which combine to regulate the total nucleotide pool of the cell. In energetically demanding tissues like skeletal and cardiac muscle, maintaining a large and tightly regulated nucleotide pool is required to keep up with the high demand of ATP consumed during muscle contraction.^7, 8^ Mutations impeding AdSS1 function are therefore expected to disrupt purinostasis and muscle energetics.

Due to its recent reporting and ultrarare nature, much of the pathophysiology of *ADSS1*-myopathy remains unknown. Animal models constitute a powerful tool for elucidating the molecular etiology of disease, and critically, for developing and testing therapeutic strategies. To date, there are two established animal models for *ADSS1* myopathy. The first is a transient zebrafish model developed with morpholino oligonucleotides to model loss of AdSS1 function.^1^ The second consists of *Adss-1* knockdown and knockout performed in *Caenorhabditis elegans* (*C. elegans)*.^3^ It should be noted that *C. elegans* possess a single *Adss* gene with only 50% sequence match to human *ADSS1*.^9^ In contrast, the human *ADSS1* gene shares 98% amino acid identity with the murine *Adss1* gene,^10^ making a mouse model an ideal candidate for disease recapitulation.

Here, we present and characterize the first mouse model of *ADSS1* myopathy, where the murine *Adss1* gene is constitutively knocked out. By examining the natural history of this mouse model, we find that homozygous *Adss1*^KO/KO^ mice are generally healthy, with subtle motor and contractility deficits. Nucleotide analysis helps shed light on the pathomechanism of this metabolic myopathy, and histological stains reveal myopathic features consistent with patient biopsy findings. Overall, we find that this mouse model recapitulates human pathology in many aspects. Importantly, this novel mouse model has quantifiable pathological features that can be used to develop and test therapies for, and further probe pathophysiology of, *ADSS1* myopathy.

## RESULTS

### Generation of constitutive *Adss1*^KO/KO^ mice

The *Adss1* knockout (C57BL/6NTac-*Adss1*^em1cyagen^) mouse line – outsourced to *Cyagen* (Santa Clara, CA, USA) for gene editing – was created by deleting exon 2 from the *Adss1* gene, resulting in a frameshift and premature termination of translation. Briefly, floxed mice were obtained by flanking exon 2 of *Adss1* with *loxP* sequences. Breeding a floxed mouse with a transgenic mouse expressing the *cre* transgene under the control of a *Zp*3 promoter ensured the excision of exon 2 in the female oocytes leading a constitutive knockout of *Adss1* in all tissues.^11^ (**Fig. 1A**). When bred to homozygosity, this germline mutation caused knockout (KO) of full-length *Adss1* in all tissues, through all stages of development.

**Fig. 1.**
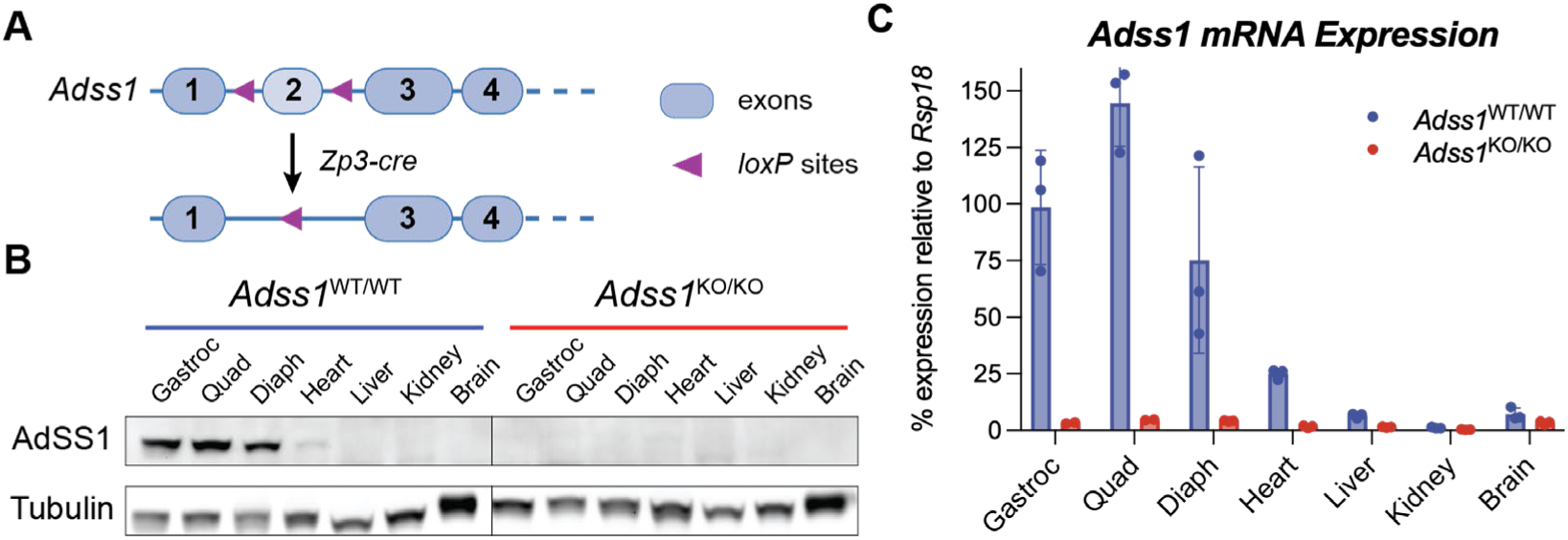
**(A)** Exon 2 of the mouse *Adss1* gene was flanked by *loxP* sites, leading to its deletion and appearance of an early stop codon upon recombination. **(B)** Western blot of tissue homogenates from *Adss1*^WT/WT^ and *Adss1*^KO/KO^ confirming the total loss of AdSS1 at the protein level in KO mice. As the muscle-specific isoform, AdSS1 is primarily expressed in striated muscle. **(C)** qRT-PCR data confirming KO of *Adss1* at the mRNA level of various tissues. Following normalization to a housekeeping gene, expression was compared to *Adss1*^WT/WT^ gastroc mRNA levels. Gastroc = gastrocnemius, Quad = quadriceps, Diaph = diaphragm.

Successful KO of *Adss1* was validated via Western blot and RT-qPCR analysis, confirming undetectable levels of AdSS1 protein and *Adss1* mRNA in various tissues of *Adss1*^KO/KO^ (or “KO” mice) compared to homozygous wild-type *Adss1*^WT/WT^ (or WT) control litter-mates (**Fig. 1B** and **C**). As expected, WT controls were confirmed to have easily detectable expression levels of *Adss1* mRNA and protein in skeletal and cardiac muscle, but not other tissue types. Expression levels of *Adss2*, the ubiquitously expressed isoform, were mostly unchanged (**SI Fig. S1**). Overall, these results demonstrate successful inactivation of the *Adss1* gene via excision of exon 2 to generate constitutive KO mice.

### *Adss1*^KO/KO^ mice are viable with no apparent weight deficit but exhibit subtle motor deficits

*Adss1*^KO/KO^ mice, along with their *Adss1*^WT/WT^ controls, were generated by crossing heterozygous *Adss1*^KO/WT^ mice with each other. Genotyping and further examination of these litters confirmed that *Adss1*^KO/KO^ mice were born in the expected Mendelian ratios and were viable. Longitudinal monitoring of body weight revealed no significant differences in the mass of *Adss1*^KO/KO^ mice compared to *Adss1*^WT/WT^ (**Fig. 2A**). In fact, *Adss1*^KO/KO^ mice appeared generally healthy, were visually indistinguishable from *Adss1*^WT/WT^, and had comparable viability up to a year (mice were not aged further). Thus, to assess possible disease phenotypes, we focused on mice that were of advanced age (10–11 months), where we hypothesized a subtle phenotype would be more evident, given the progressive nature of the disease in humans. We conducted a series of functional assessments to determine if *Adss1*^KO/KO^ mice exhibited any pathological neuromuscular phenotypes, including defects in running abilities or functional strength.

**Fig. 2.**
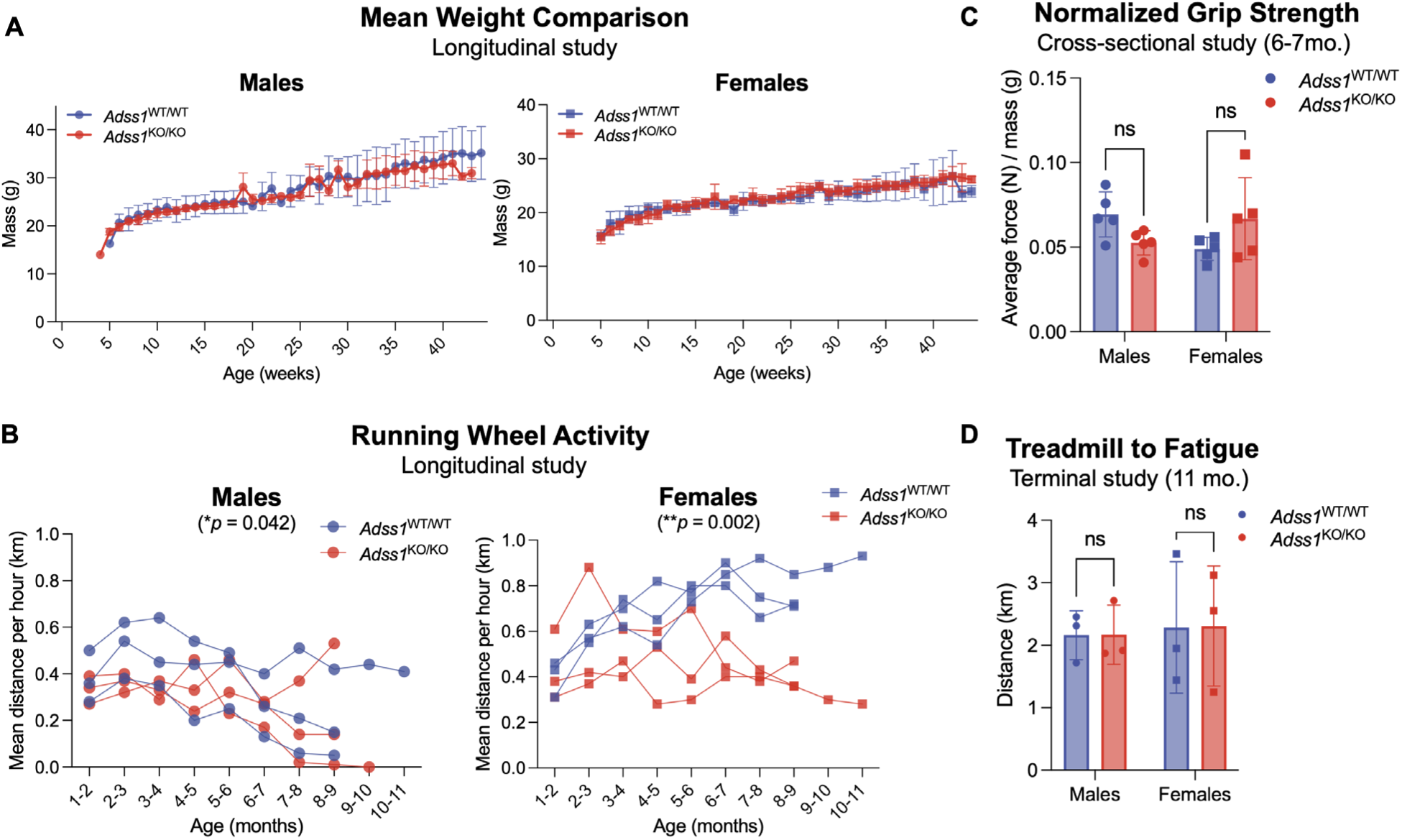
**(A)** Longitudinal analysis of mean weight between *Adss1*^KO/KO^ and *Adss1*^WT/WT^ mice (n = 5 per group). **(B)** No significant difference in grip strength between *Adss1*^KO/KO^ and *Adss1*^WT/WT^ mice, accounting for sex-based differences in weight (n = 5 per group). **(C)** Small difference observed in voluntary running distance between *Adss1*^KO/KO^ and *Adss1*^WT/WT^ mice, based on spontaneous activity (n = 3 per group). **(D)** No significant difference in endurance between *Adss1*^KO/KO^ and *Adss1*^WT/WT^ mice when ran to exhaustion on a treadmill (n = 3 per group).

To monitor running abilities longitudinally, voluntary running activity was recorded with in-cage running wheels for approximately 11 months across 12 mice. Mice were singly housed, and each running wheel tracked the number of wheel rotations (equivalent to distance run) per hour. Within this longitudinal study, we observed a difference in running distance between *Adss1*^KO/KO^ and *Adss1*^WT/WT^ mice (\**p* = 0.04 in males, \*\**p* = 0.002 in females; **Fig. 2B**). Interestingly, *Adss1*^KO/KO^ females exhibited a larger difference in running distance than males, with a divergence from wild-type values beginning around 6–7 months of age.

Given the subtle difference emerging, we studied four limb grip strength as a non-terminal metric of muscle function at 6–7 months of age. Overall, we observed no difference in the grip strength of *Adss1*^KO/KO^ versus *Adss1*^WT/WT^ mice, in either sex (**Fig. 2C**).

Beyond voluntary running and grip strength, we aged additional mice (who were not given running wheels) to 11 months of age to assess the effect of *Adss1* loss-of-function on running endurance. These 12 mice were subjected to forced uphill running on a treadmill programmed with increasing speed over time as a terminal cross-sectional study. Ultimately, the fatigue-induced treadmill test did not indicate a difference in endurance threshold between *Adss1*^KO/KO^ and *Adss1*^WT/WT^ mice, in either sex, when run to exhaustion (**Fig. 2D).** Based on our findings from these three motor function tests, *Adss1*^KO/KO^ mice appear generally healthy and perform at comparable levels to *Adss1*^WT/WT^ mice, with only subtle differences in voluntary running distance.

### *Adss1*^KO/KO^ mice exhibit muscle contractile defects when examined *ex vivo*

Because *Adss1*^KO/KO^ mice exhibited only subtle motor deficits in cross-sectional and longitudinal activity studies, we elected to examine their muscle function more closely using *ex vivo* contractile force and fatigue measurements. We studied the soleus and extensor digitorum longus (EDL) of 3- and 10-month-old mice as the EDL is exclusively composed of glycolytic fast twitch fibers while the soleus is the only hindlimb muscle in the mouse with substantial populations of oxidative fibers (roughly 50% type I and 50% type IIA).^12^ Therefore, these two muscles represent extremes in murine muscle metabolism, and are small and spindle-shaped, making them optimal for *ex vivo* contractility measurements. Similarly to whole body weight, there was no significant difference in the mass of the EDL and soleus of *Adss1*^KO/KO^ mice when compared to their WT counterparts (**SI Fig. S2**). However, in response to supramaximal stimulation, *Adss1*^KO/KO^ EDL generated lower peak contractile force than *Adss1*^WT/WT^ counterparts (**Fig. 3A**), despite being of a comparable size. Contractility differences across genotypes were not significant for the soleus, although the peak contractile force similarly trended lower in *Adss1*^KO/KO^ mice.

**Fig. 3.**
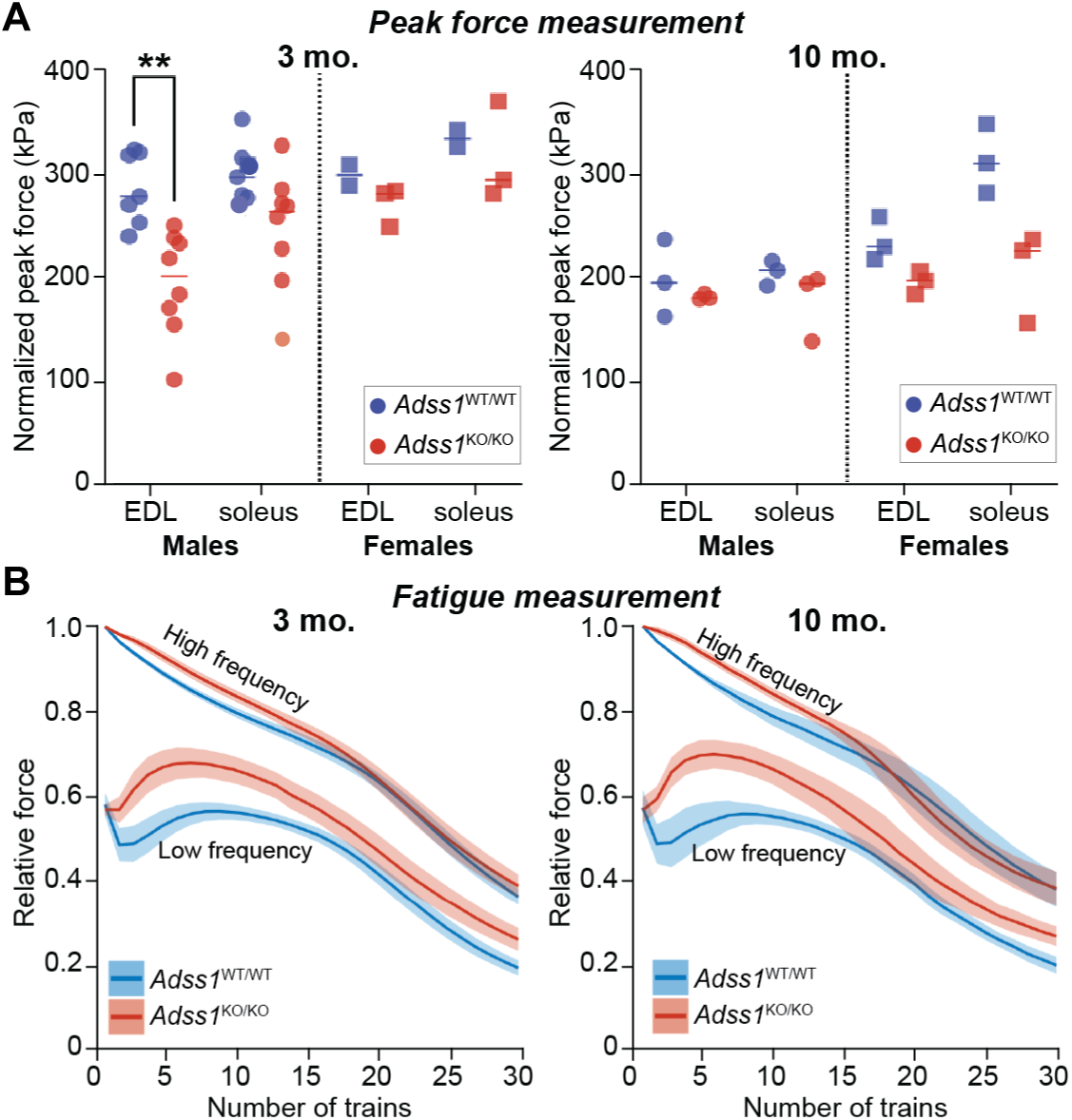
**(A)** Comparison of cross-sectional area-normalized peak force generated by muscles in response to electrical stimulation, of 3- and 10-month-old mice. A statistically significant difference in contractility was observed in the EDL of male mice (n = 7 for *Adss1*^WT/WT^ and 8 for *Adss1*^KO/KO^) with a similar (albeit not significant) trend in other groups. **(B)** *Adss1*^KO/KO^ EDLs react differently to a low frequency fatigue protocol. While the force exerted by *Adss1*^WT/WT^ initially decreases upon repeated low frequency stimulation, the force of *Adss1*^KO/KO^ increases, suggesting a possible relaxation defect. Force measured is normalized to the initial force produced following high frequency stimulation. Mean value is plotted, with ribbons representing the 95% confidence interval.

We then examined susceptibility to fatigue in a subsequent *ex vivo* analysis measuring the relative force generated from a stimulus train consisting of a high frequency component that produced peak tetanic force, and a low frequency component that elicited roughly 60% of tetanic force in the rested muscle (**SI Fig. S3**). During the first 10 high frequency contractions, *Adss1*^KO/KO^ EDLs generated slightly lower relative high frequency force than their WT counterparts, but from the 15^th^ to 30^th^ trains of stimulation, this difference subsided. Interestingly, at low frequency, the first five trains of stimulation resulted in a higher force generation in the *Adss1*^KO/KO^ EDLs that reached nearly 70% of the initial force at high frequency compared to *Adss1*^WT/WT^ EDLs which generated a force that was <60% of the initial force (**Fig. 3B**). In other words, at a number of repeated low frequency pulses, the force generated by *Adss1*^KO/KO^ EDLs initially increased before steadily declining. Again, this difference was only observed in the EDL of *Adss1*^KO/KO^ mice and not the soleus (**SI Fig. S4**).

### Loss of AdSS1 is associated with a reduced total nucleotide pool without gross dysregulation of purinostasis

To get a better glimpse into skeletal muscle energetics in response to *Adss1* KO, we measured the concentration of nucleotides in various muscles of wild-type and *Adss1*^KO/KO^ mice. We examined nucleotide levels in both rested muscles, harvested from anesthetized mice, and fatigued muscles, measured following the *ex vivo* fatigue protocol. Overall, *Adss1*^KO/KO^ muscles contained significantly lower concentrations of total nucleotides when compared to wild-type, in both rested and fatigued muscle (**Fig. 4A**). Interestingly, the [ATP]/[ADP] ratio – a proxy for available free energy^13^ – was mostly maintained between muscles of the two genotypes (**Fig. 4B**), indicating that despite a smaller nucleotide pool, muscle energetics and purinostasis were not globally dysregulated by loss of *Adss1*. Consistent with AdSS1 enzymatic dysfunction, the concentration of IMP – its substrate – was significantly higher in rested *Adss1*^KO/KO^ muscles compared to rested *Adss1*^WT/WT^ muscles. After intense contractions and profound fatigue leading to degradation of adenine nucleotides, IMP concentrations rose in both KO and WT muscle, but consistent with a shrunken nucleotide pool, the IMP content of *Adss1*^KO/KO^ muscles was significantly lower compared to *Adss1*^WT/WT^ muscles once fatigued (**Fig. 4C**).

**Fig. 4.**
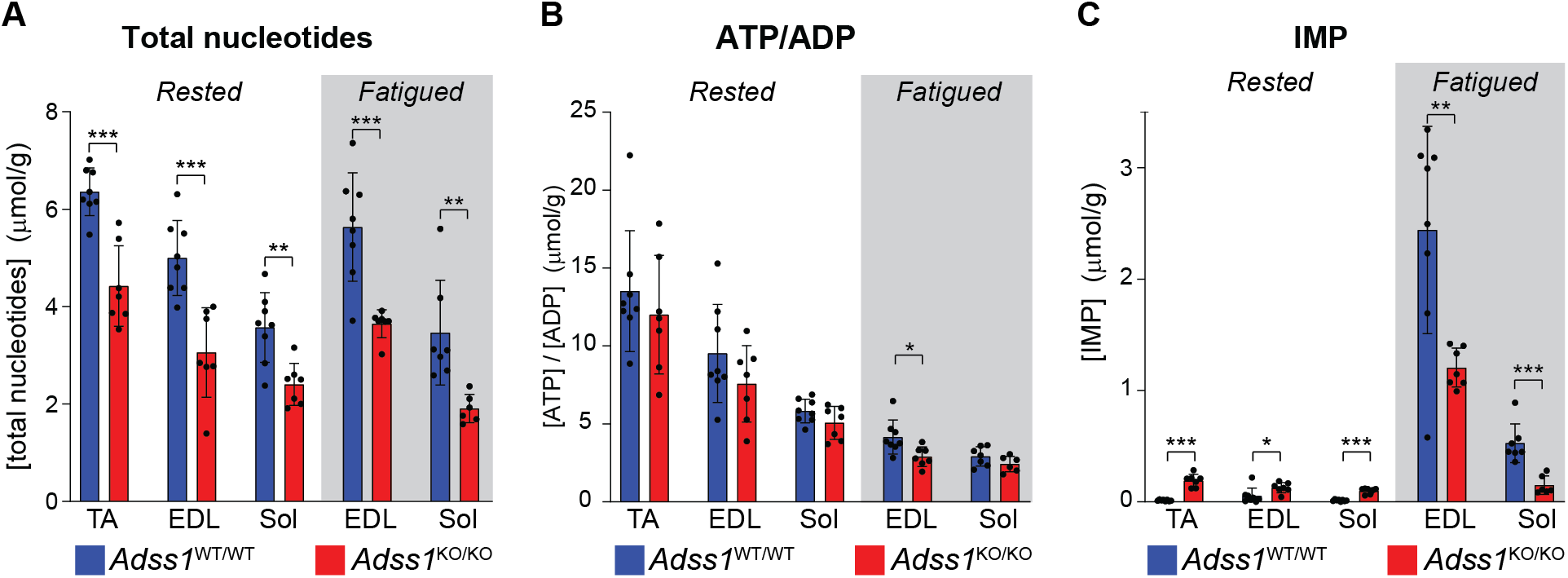
Concentration of various nucleotides in muscle homogenate from *Adss1*^WT/WT^ and *Adss1*^KO/KO^ mice, in rested and fatigued muscles. **(A)** Total nucleotide concentrations were significantly lower in all *Adss1*^KO/KO^ muscles compared to *Adss1*^WT/WT^ muscles. **(B)** ATP to ADP ratios remained mostly unchanged between muscles of *Adss1*^KO/KO^ and *Adss1*^WT/WT^ mice, in both rested and fatigued states. **(C)** IMP levels were significantly higher in rested *Adss1*^KO/KO^ muscles, consistent with an *Adss1* bottleneck. Once fatigued, IMP concentrations rose in both KO and WT muscle, but consistent with a smaller nucleotide pool, the IMP content of fatigued *Adss1*^KO/KO^ muscles was significantly reduced compared to fatigued *Adss1*^WT/WT^. TA: tibialis anterior, EDL: extensor digitorum longus, Sol: soleus. (\**p* < 0.05; \*\**p* < 0.01; \*\*\**p* < 0.001 as assessed by unpaired *t* test.)

### AdSS1 deficient muscles show marked myopathic features by histology

To assess whether *Adss1*^KO/KO^ mice displayed any of the myopathic features identified in patient biopsies, tibialis anterior (TA) muscle cross-sections from the cohort of 20 aged (11-month-old) mice were stained with modified Gömöri Trichrome. Several myopathic features (rimmed vacuoles, fiber splitting, or internal nuclei) were apparent in all *Adss1*^KO/KO^ mice (**Fig. 5A and B**, and **SI Fig. S5**). Notably, *Adss1*^KO/KO^ mice were found to have increased numbers of features resembling rimmed vacuoles^2, 14, 15^ compared to wild-type controls, with a clear sex-based difference. By 11 months of age, rimmed vacuoles were observed in 12% of male *Adss1*^KO/KO^ TA fibers, versus 3% in male *Adss1*^WT/WT^ fibers (*p* = 0.004, n = 5). As for 11-month-old females, about 6% of *Adss1*^KO/KO^ TA fibers contained rimmed vacuoles, compared to only 0.5% of female *Adss1*^WT/WT^ fibers (*p* = 0.004, n = 5) (**Fig. 5B**).

**Fig. 5.**
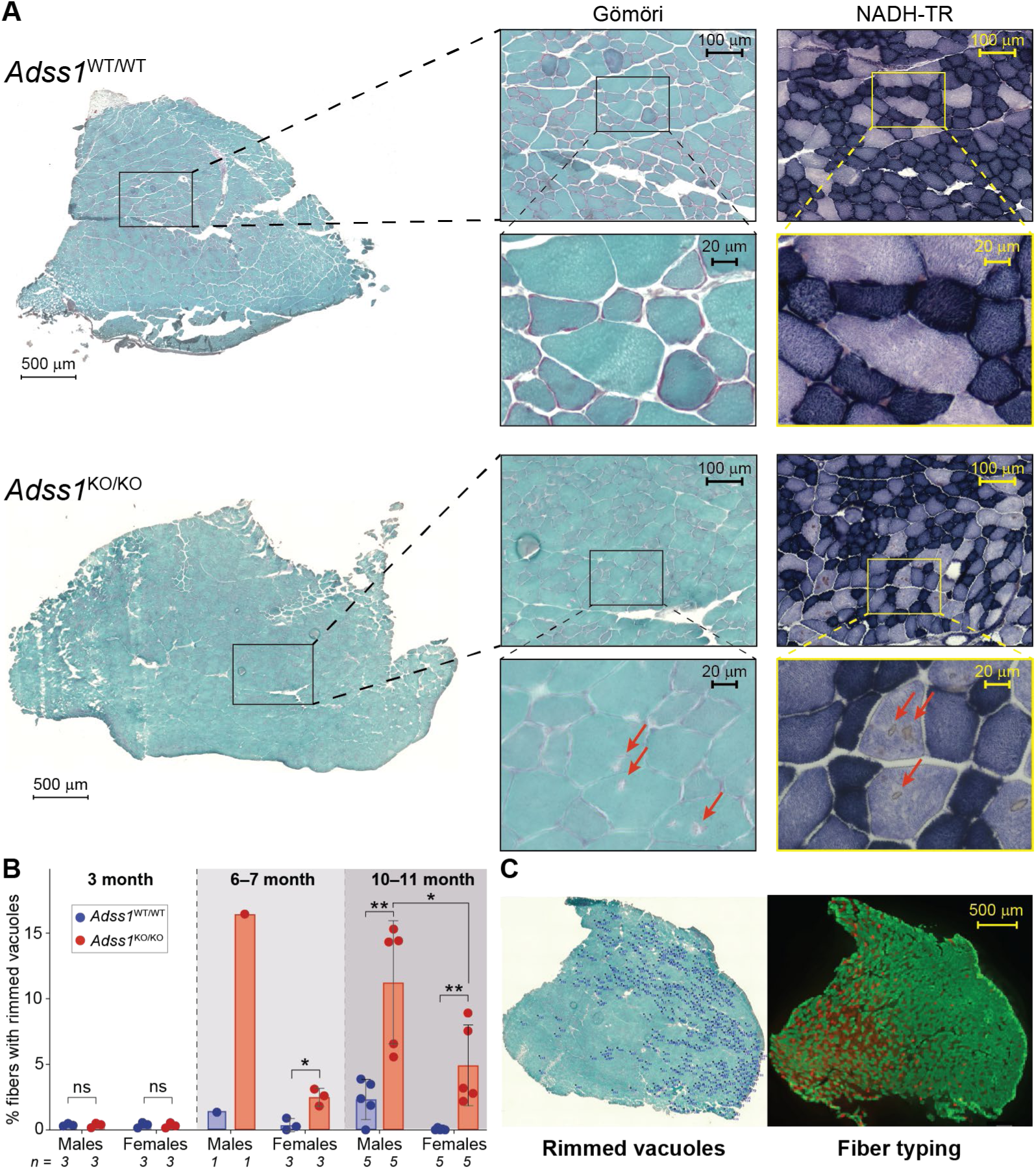
**(A)** Modified Gömöri Trichrome and NADH staining on serial sections of *Adss1^WT^*^/WT^ and *Adss1*^KO/KO^ TA (10.5 mo.). **(B)** Proportion of rimmed vacuole-positive fibers in TA muscle of 40 mice quantified from Gömöri Trichrome staining. Significant differences between *Adss1*^KO/KO^ and *Adss1^WT^*^/WT^ mice beginning at 6 months of age, with a larger difference observed at 10–11 months. At 10–11 months, there was a sex-based difference in rimmed vacuole positive fibers, with *Adss1*^KO/KO^ males having - 6.4 ± 2.6% more rimmed vacuole-positive fibers than *Adss1*^KO/KO^ females. (\**p* < 0.05; \*\**p* < 0.01; \*\*\**p* < 0.001) **(C)** Parallel processing of modified Gömöri Trichrome (left) and immunofluorescent fiber type staining (right) of *Adss1*^KO/KO^ TA at 10.5 months old. Distribution of myofibers with rimmed vacuoles (dark blue tags) aligned with type IIB glycolytic fibers (green) from fiber type staining.

To better characterize the location of the damage, additional serial sections were stained with NADH-TR to visualize oxidative activity modulated by tetrazolium reductase, with high oxidative activity accumulating an indigo hue. Here we found rimmed vacuoles manifesting as focal brown puncta in fibers with least oxidative activity (pale purple), (**Fig. 5A**). To further validate these findings, we compared serial sections stained with Gömöri Trichrome to immunofluorescently fiber typed sections. While we did not observe a difference in fiber type composition between *Adss1*^KO/KO^ muscles and wild-type controls (**SI Fig. S6**), *Adss1*^KO/KO^ mice were found to have more fiber damage in type IIB glycolytic fibers (in green), as evidenced by the localization of rimmed vacuoles in parallel fiber type staining (**Fig. 5C** and **SI Fig. S7**).

### AdSS1 deficient muscles exhibit abnormal glycogen distribution

Upon visualizing fiber damage with both NADH-TR and Gömöri Trichrome stains, we conducted additional serial stains to interrogate the composition of the damaged fiber areas. Based on comparable histological markers in Pompe disease, in which rimmed vacuoles are associated with abnormal glycogen accumulation,^16^ we stained additional sections from *Adss1*^KO/KO^ TA muscle with periodic acid-Schiff (PAS) to visualize glycogen. Interestingly, sections of *Adss1*^KO/KO^ mice revealed that glycogenic components (glycogen, glycoproteins, glycolipids) were similarly accumulating in fibers (**Fig. 6A**).

**Fig. 6.**
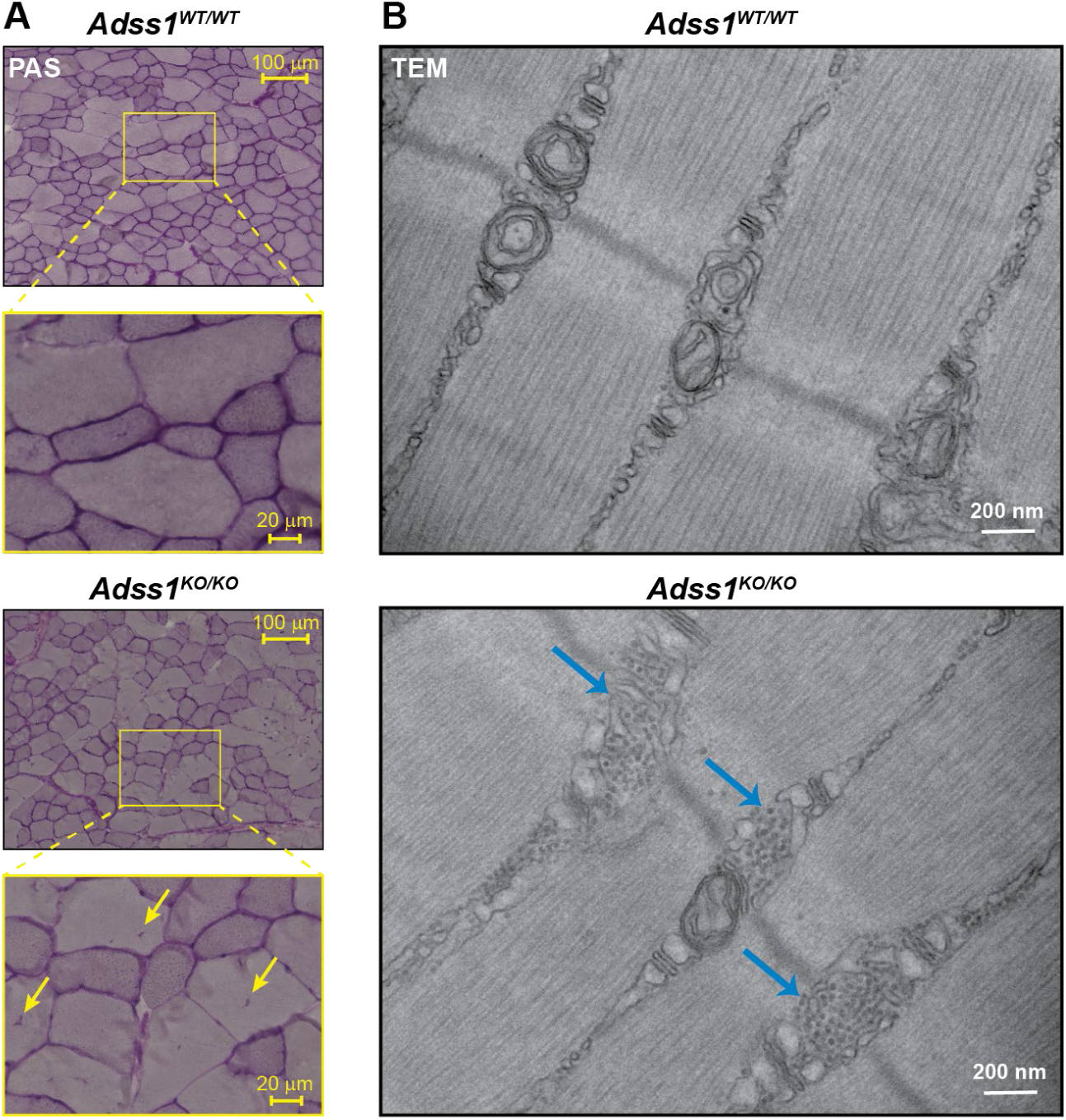
**(A)** PAS staining on sections highlighted accumulation of glycogenic components in cross-sections of *Adss1*^KO/KO^ muscle (yellow arrows). **(B)** At higher magnification, TEM revealed glycogen aggregates in longitudinal sections of *Adss1*^KO/KO^ muscle, identifiable as small dark circles, highlighted by the blue arrows. These aggregates were largely absent in wild-type controls.

To further investigate the ultrastructural changes associated with AdSS1 deficiency, transmission electron microscopy (TEM) was performed on longitudinal sections of the TA from *Adss1*^KO/KO^ mice and wild-type controls. TEM analysis revealed prominent glycogen deposits localized near the T-tubules in *Adss1* KO muscles, a pattern that was only occasionally observed in controls (**Fig. 6B**). These dense glycogen aggregates (designated by blue arrows) were situated close to key sites of muscle excitation-contraction coupling. In both WT and *Adss1* KO muscles, mitochondria were typically found in areas flanking the Z-lines; however, in *Adss1* KO muscles, mitochondria seemed smaller or absent in many areas where glycogen was accumulated. While examined at a vastly different scale and orientations, the observation of numerous glycogen aggregates by TEM supports the histopathological evidence of abnormal glycogen accumulation and suggests a possible disruption of glycogen homeostasis due to loss of AdSS1.

## DISCUSSION

In this work, we developed and characterized a novel mouse model of *ADSS1* myopathy. Longitudinal studies revealed that these constitutive *Adss1*^KO/KO^ mice were generally healthy with no weight differences and only minimal motor function defects when compared to wild-type mice. *Adss1*^KO/KO^ mice exhibited comparable endurance and grip strength, and only differed in voluntary running distance when aged, although this difference was minimal in males and of larger magnitude in females. More notable pathological phenotypes were observed by *ex vivo* contractility and fatigue studies, nucleotide content measurements, and histopathological and TEM analyses.

While physiologic differences in contractility and fatigue were not apparent by grip strength and endurance measurements, when examined *ex vivo*, *Adss1*^KO/KO^ EDLs produced significantly less force compared to *Adss1*^WT/WT^ EDLs, despite their comparable size. Moreover, *Adss1*^KO/KO^ EDLs exhibited a distinct response to a low-frequency fatigue protocol. The initial increase in force observed during repeated low-frequency stimulation may indicate a defect in muscle relaxation: if the muscle fails to fully relax between stimuli, force output can compound even if the force produced by each individual stimulus is reduced. This difference was evident in the EDL but not in the soleus muscle of *Adss1*^KO/KO^ mice, likely due to differences in their fibers’ dominate metabolic pathways. Specifically, the EDL is composed almost exclusively of glycolytic fibers (type IIB and IIX), which are more prone to fatigue compared to the soleus, in which oxidative fiber types (I and IIA) make up about 85% of muscle fibers.^12, 17^

Metabolically, examination of various nucleotide concentrations in muscle offered novel insight into the pathomechanism of *ADSS1* myopathy. *Adss1* KO caused a reduction in the total nucleotide pool without globally dysregulating purinostasis. This finding agrees with the notion that ATP/ADP ratio can be regulated independently from ATP levels.^18^ ATP levels are regulated by the *de novo* purine synthesis and the purine salvage pathways, both of which involve AdSS. The IMP to s-AMP conversion that AdSS1 catalyzes is the canonical pathway for newly synthesized purines to enter the adenine nucleotide pool in muscles.^19, 20^ Detection of ATP nucleotides in *Adss1*^KO/KO^ muscles – albeit at lower levels – and the viability of the mice suggest a form of metabolic compensation that may account for the mild physiological deficits of motor function in mice. One likely possibility for compensation is *Adss2*, the non-muscle isoform of this enzyme, which is ubiquitously expressed at low levels and catalyzes the same reaction, although with slightly different kinetic parameters.^21^ Interestingly, in preliminary work, we did not observe an upregulation of *Adss2* mRNA in *Adss1*^KO/KO^ tissues (**SI Fig. S1**). It will be interesting to further investigate this potential compensatory mechanism in future work and to examine whether evidence for similar compensation is absent in human tissues from patients with frank weakness.

Histologically, we found myopathic features in *Adss1*^KO/KO^ mice that correlate well with observations from patient findings, particularly for fiber splitting, internal nuclei, and rimmed vacuoles.^1, 2, 14, 15, 22^ In fact, the most common myopathic feature we observed in *Adss1*^KO/KO^ mice were rimmed vacuoles, which were commonly observed by 6 months of age. This aligns with quantifications from patient biopsies which reported rimmed vacuoles in about 2% of total fibers in cohorts of Japanese and Korean patient.^1, 2, 14, 15, 22, 23^ Most *Adss1*^KO/KO^ mice were found to have far more rimmed vacuole-positive fibers than patients (average of 11% and 5% for 11-month-old *Adss1*^KO/KO^ male and female mice, respectively, **Fig. 5B**). This discrepancy is not unusual given that mice have different fiber type makeup than humans, including type IIB fibers (extreme fast-twitch, glycolytic) that do not exist in humans.^24^ Despite manifesting only subtle neuromuscular phenotypes and appearing generally healthy (**Fig. 2**), *Adss1*^KO/KO^ mice recapitulate the disease phenotype found in patients at a histological level (**Fig. 5**).

A few of our measurements revealed sex differences, although it is not apparent that mice of one sex are more severely affected than the other. Notably, we observed a decrease in the voluntary running distance of older female *Adss1*^KO/KO^ mice compared to wild-type, but not in males. Conversely, histopathological analyses revealed that male *Adss1*^KO/KO^ mice had considerably more myopathic features compared to female *Adss1*^KO/KO^ mice starting at 6 months of age, as did wild-type males compared to wild-type females. So far, no sex differences have been noted in *ADSS1* myopathy patients, although this may be a product of the small number of cases reported to date.

Perhaps the most notable finding of this work is a striking fiber type-specific involvement in pathology. *Ex vivo* studies revealed greater functional impairment in the highly glycolytic EDL compared to the more oxidative soleus. Consistent with this finding, histological analyses underscored that fiber damage in *Adss1*^KO/KO^ muscle was concentrated in type IIB fibers. This fiber-type specificity supports a metabolic basis for fiber damage: since type IIB fibers are the most glycolytic and fatigue-prone, the observed vulnerability may stem from the metabolic properties specific to this fiber type in the context of AdSS1 deficiency. Indeed, fast fibers generally have higher levels of adenine nucleotides^25^ (**Fig. 4**, comparing total nucleotides in EDL vs soleus) and may therefore be more sensitive to loss of nucleotides. Taken together, these data reveal that loss of AdSS1 preferentially affects glycolytic fiber populations.

Beyond fiber-type specificity, ultrastructural imaging revealed abnormalities in glycogen distribution, reinforcing the idea that glycolytic pathways may be centrally involved in the pathophysiology of the disease.^26^ These findings suggest that, although AdSS1 deficiency is *a priori* a disruption of purine nucleotide metabolism, it may also trigger downstream disturbances in glycogen handling, representing a possible convergent pathological mechanism with glycogen storage diseases, such as Pompe disease.^27^ While enzyme replacement therapy has substantially improved cardiac outcomes in Pompe, it has not fully corrected glycogen mislocalization or the associated skeletal myopathy.^28^ This shortcoming underscores the possibility that, although likely secondary to the disruption of purinostasis, these glycogen abnormalities may need to be particularly addressed to achieve complete therapeutic benefit in *ADSS1* myopathy.

Overall, this *Adss1*^KO/KO^ mouse represents the first mammalian model of *ADSS1* myopathy, addressing key limitations of previous systems. Unlike *C. elegans*, which lacks a second isoforms of AdSS,^9^ and transient zebrafish morpholino models,^1^ this stable knockout in mice offers a more genetically and metabolically relevant platform for studying the disease. Importantly, mice share high genetic and metabolic similarity with humans and express both known *ADSS1* isoforms,^6, 10^ rendering this model well-suited for translational research. While these mice appear largely healthy under unstressed conditions, they do recapitulate important aspects of the human disease, including muscle weakness (when examined *ex vivo*) and hallmark histological features. Future studies using physiological stressors or sensitized genetic backgrounds may help unmask more severe phenotypes and better model the progressive aspects of *ADSS1* myopathy.

In summary, the *Adss1*^KO/KO^ mouse provides a promising tool for dissecting the pathophysiology of *ADSS1* myopathy and serves as a valuable preclinical platform for evaluating potential therapies. The observed fiber type-specific pathology offers new insight into disease mechanisms and moves us closer to understanding and treating this rare disease.

## METHODS

### Animal Conditions

All mice were housed in the Animal Research Core Facility (ARCH) at Boston Children’s Hospital in accordance with the Institutional Animal Care and Use Committee (IACUC). Mice were maintained with a standard light/dark cycle, given standard chow, a grain-based rodent diet in pelleted form, and accessed water via lixit.

### *Adss1*^KO^ Design

The *Adss1*^KO^ line, outsourced to Cyagen for gene editing, was generated by introducing *loxP* sites flanking exon 2 of Adss1 (C57BL/6N-Adssl1^em1cyagen^, serial number CKOCMP-11565-Adssl1-B6N-VA). Mice were bred by Charles River Laboratories on a C57BL/6N background and sent to Cyagen for Cre-lox manipulation and CRISPR gene editing to create the *Adss1* KO models. Homozygous *Adss1* floxed mice were crossed with mice expressing the Cre recombinase transgene under a Zp3 (zona pellucida 3) promoter on a C57BL/6J background. The heterozygous *Adss1*^KO/WT^, Cre positive mice were shipped to the Beggs lab, where they were further crossed to wild-type C57BL/6J mice to remove the Cre transgene. Experimental mice were generated by crossing heterozygous (and Cre negative) *Adss1*^KO/WT^ mice in order to compare *Adss1*^KO/KO^ and *Adss1*^WT/WT^ littermates.

### Primers for Genotyping

Mice were genotyped using the following primers using a touchdown PCR protocol with short elongation time (15 sec):

*Adss1*-F1: 5’-AATTCCCAAAAGAAACAGCCTCC-3’

*Adss1*-F2: 5’-CCTGGAGCCCCCAGCCAGAA-3’

*Adss1*-R: 5’-TTCTTCATCAACCAGAGAGACCAG-3’

Primers F1 + R amplify a 1045 bp amplicon in the wild-type *Adss1* gene and a 187 bp amplicon in the knockout gene. Furthermore, primers F2 + R amplify a 357 bp amplicon in the wild-type gene, but no amplicon (F2 does not bind) in the knockout gene.

### Western Blot Analysis

Various tissues were extracted from euthanized mice, frozen, and stored at −80°C until processing. For muscle tissues, samples were frozen in liquid nitrogen-cooled isopentane to preserve morphology for possible downstream histological analyses. For Western blot analysis, a small portion of tissue was collected, and added to T-PER tissue protein reagent (Thermo Fisher Scientific) supplemented with an EDTA-free protease inhibitor tablet (Roche). Samples were homogenized using 3.2 mm stainless steel RNAse-free milling beads (Next Advance) and a bullet blender (Next Advance) until no large tissue pieces remained. Homogenates were centrifuged in a tabletop centrifuge at max speed for 5 min at 4°C to pellet tissue debris. Supernatant was collected and protein content was quantified using a BCA assay (Pierce) according to manufacturer’s instructions. Following normalization, samples were briefly boiled in 1× sample loading dye (Novex) and loaded on a precast 4–12% NuPAGE Tris-Acetate gel (Invitrogen) and wet-transferred to a PDVF blot. Ponceau stain (Millipore) was used to assess even transfer. The blot was blocked in Everyblot blocking buffer (Bio-Rad), then probed for AdSS1 using a custom rabbit polyclonal peptide antibody (Biomatik, raised against the SGTRASNDRPPGAGGVKRGR-Cys peptide) (1:500 dilution in Everyblot), followed by a secondary goat anti-rabbit Starbright Blue 700 (Bio-Rad, 1:5,000 dilution) and a rhodamine-labeled anti-tubulin antibody (Bio-Rad, 1:5,000 dilution) as a loading control. Blots were washed then imaged using a ChemiDoc MP Imaging System (Bio-Rad).

### RT-qPCR Analysis

For RT-qPCR analysis, tissues were collected from 3 *Adss1*^WT/WT^ and 3 *Adss1*^KO/KO^ mice, which served as biological replicates. RNA from each tissue was analyzed 3 times, serving as technical replicates. Total RNA was extracted from samples by homogenizing various murine tissues in TRIzol (Thermo Fisher Scientific), using the beads and bullet blender described above (Next Advance). Following a chloroform extraction, the aqueous supernatant was collected and purified using a Direct-zol RNA kit (Zymo) according to manufacturer’s instructions, including the optional DNAse treatment. RNA concentrations were measured by Nanodrop. Following normalization, real time quantitative PCR was performed using the Reliance One-step Multiplex RT-qPCR MasterMix (Bio-Rad) which combines cDNA synthesis and RT-qPCR steps in a single mix. TaqMan-based chemistry probes (Applied Biosystems) were designed to specifically quantify *Adss1* (FAM) (Applied Biosystems, 4351370), *Adss2* (VIC) (Applied Biosystems, 4448490), and *Rsp18* (Tex615) (Biorad, qMmuCEP0053856) RNA levels using an ABI Prism 7300 Real Time PCR System. Gene expression for each sample was normalized using *Rsp18* as a housekeeping gene, then technical replicate values were averaged. Relative expression was calculated compared to *Adss1*^WT/WT^ gastrocnemius samples.

### Voluntary Running Wheels

Longitudinal studies were performed on a set of 12 mice (6 *Adss1*^WT/WT^, 6 *Adss1*^KO/KO^). To assess voluntary running distance, each mouse was individually housed with a low-profile wireless running wheel (Med Associates Inc.) starting at 4 weeks old that wirelessly tracked their total distance ran per day. Running wheel data was received automatically via Bluetooth connection to a local computer using the Wheel Counter Utility software (Med Associates Inc.) and exported periodically. Once mice reached 11 months, running wheels were removed and all data was compiled.

### Grip Strength

At 6–7 months of age, a basal grip strength test was performed on the longitudinal cohort along with 8 other sex- and age-matched mice (for a total of 20 mice). The BIO-GS3 grip strength meter from BioSeb assesses mouse neuromuscular function by measuring the peak force (N) it generates after being tugged by the tail. The maximal peak force (N) is reflective of its basal muscular grip strength in their forelimbs and hindlimbs. Two measurements were taken for each mouse as technical replicates to account for any errors in handling the mouse.

### Treadmill to Exhaustion

Additional mice were aged to ∼11 months and subjected to a premortem treadmill test where they were run to exhaustion on an automated single belt rodent treadmill (Maze Engineers). These mice were not on running wheels at any point in their life, so as to measure basal (not training-based) endurance. A treadmill protocol adapted from Castro & Kuang’s study^29^ was used to assess endurance threshold with a progressive increase in speed. The treadmill was positioned at an 11° incline with mice running uphill. Initial speed was set to 3.3 m/min for 5 min with no shock to allow the mice to acclimate to the treadmill. Subsequent speed progression was as follows: 8 m/min for 5 minutes, 10 m/min for 10 minutes, 12 m/min for 20 minutes, 15 m/min for 20 minutes, and 18 m/min for 20 minutes, with a maximum speed reaching 25 m/min. The point of exhaustion was determined to be a cessation of running for five seconds and no attempt to continue running after electrical shock.

### *Ex vivo* Peak Force Measurements

Mice were anesthetized by an intraperitoneal injection of pentobarbital (80 mg/kg) and supplemented with 100% O_2_ delivered through a nose cone. One soleus and one EDL were carefully removed and submerged in room temperature bicarbonate buffer equilibrated with 95% O_2_, 5% CO_2_. Silk treads were tied to each tendon. The muscles were then transferred into a bath containing gas equilibrated bicarbonate buffer (95% O_2_, 5% CO_2_) maintained at 35°C. The silk treads were used to attach the muscles between a muscle lever system (Aurora Scientific) and a fixed post. Platinum stimulating electrodes flanked each muscle, as previously described.^30^ Brief tetnanic contractions were used to determine the optimal muscle length for tension generation. Peak force generated by each muscle was compared between genotypes, after normalization to the cross-sectional area of the respective muscles (as described in the following section).

### *Ex vivo* Force-Frequency Relationship and Muscle Fatigue

Each muscle was subjected to fixed-end contractions at stimulation frequencies eliciting twitches up to fully fused tetani. These force-frequency data were fit by a sigmoid curve.^30^ The inflection point of each curve, which elicits roughly 60% of tetanic force, was used to determine the stimulation frequency for eliciting the muscle’s “low frequency” force. “High frequency” force was the frequency used to elicit peak tetanic tension. Each muscle was subsequently subjected to 30 stimulus trains consisting of a low-frequency stimulation phase and a high-frequency stimulation phase. Trains were separated by 3 seconds.

Immediately after the final train, the muscle was removed from the bath, the tendons trimmed, and the muscles quickly weighed. Muscles were then immediately frozen by freeze-clamping and used for purine analysis.

### Normalization of Muscle Force

Muscle mass, optimal muscle length, and published muscle length to fiber length ratios were used to estimate muscle physiological cross-sectional area, which was subsequently used to normalize muscle force.^30^

### Purine Measurements

For ‘rested’ measurements, one TA, one EDL, and one soleus muscles were removed from mice while anesthetized with pentobarbital. For ‘fatigued’ measurements, the contralateral EDL and soleus were collected promptly after *ex vivo* fatigue protocol. Muscles were freeze-clamped immediately. Metabolites were extracted in 50-fold (volume : muscle weight) ice-cold 80% ethanol with rapid homogenization in glass-on-glass tissue grinders (Kontes). Extracts were centrifuged at 4°C to remove precipitate. Samples were stored at −80°C until analysis.

Concentrations of adenine nucleotides (ATP, ADP, and AMP), and related products (IMP, adenosine, adenine, and inosine) were determined by ultra performance liquid chromatography (UPLC) using a Waters Acquity Premier UPLC H-class system as described previously.^31^ Briefly, separation was achieved by gradient reverse-phase chromatography using an Acquity UPLC HSS T3 1.8 μm, 2.1 × 150 mm column (Waters). Metabolites were identified by comparison of 254nm peak retention times to those of commercially available standards (Sigma-Aldrich) and confirmed by mass detection (Acquity QDa, Waters).

### Histology

#### Tissue Preparation for Histology

For all histological stains, mouse tibialis anterior (TA) muscles were harvested and flash frozen in liquid nitrogen cooled isopentane for homogenous freezing and stored at –80°C until cryosectioning. TA muscles were selected for their size and varied fiber type composition. Frozen muscles were embedded in *Tissue-Tek®* (OCT) compound and clamped with dry ice to preserve vertical orientation of fibers. Sections were cut at 10 micrometers and adsorbed to *SuperFrost*^TM^ *Plus* slides to promote tissue adhesion. At least 3 mice were used for each experimental group (age and sex-matched), at 3 months, 6–7 months and 10–11 months, and stained.

#### Histological Staining

All sections were stained serially with hematoxylin-eosin (H&E) for morphological analysis of muscle, modified Gömöri Trichrome to identify myopathic features, nicotinamide-adenine dinucleotide tetrazolium reductase (NADH-TR) to visualize skeletal muscle fiber types based on oxidative activity, immunofluorescent fiber type staining to visualize myosin heavy chain subtypes (type 1A, 2A, 2B, 2X) based on fluorophore conjugated antibodies, and periodic acid-Schiff (PAS) to label glycogen aggregates. All slides were stored at -80°C prior to use and airdried for approximately 20 minutes before each staining procedure.

H&E staining was performed with an initial 5-minute fixation in cold methanol, 5-minute wash in phosphate buffered saline (PBS), 7-minute counterstain in hematoxylin, quick rinses in deionized water followed by acid alcohol, deionized water, 1% ammonia water, and deionized water before staining for 1 minute in eosin. Slides were dehydrated in a series of 70%, 95% and 100% ethanol and xylenes before mounting with Cytoseal (Avantor) with a coverslip.

Modified Gömöri Trichrome staining was performed with an initial 2-minute rehydration in deionized water prior to hematoxylin counterstaining for 5 minutes to visualize nuclei and rimmed vacuoles. The rehydration step was critical to ensure that fibers stain turquoise without diffuse purple staining artifacts across the tissue. Following a 5-minute wash in running tap water, slides were stained in Gömöri Trichrome One-Step solution (Newcomer Supply) for 15 minutes and then washed in running tap water for 2 minutes before differentiating in 0.5% acetic acid and dehydrating in a series of ethanol and xylenes. Internal nuclei were visualized as punctate purple dots with hematoxylin and rimmed vacuoles were visualized as damaged fibers with vacuoles rimmed with a magenta hue.

For fiber type staining via oxidative activity modulated by nicotinamide-adenine dinucleotide tetrazolium reductase (NADH-TR), slides were incubated in an NADH solution (0.16% NADH in 50mM Tris-HCL at pH 7.6) with Nitro-Blue Tetrazolium for 2 hours at 37°C to form a blue formazan precipitate at the site of oxidative activity. Darker fibers indicated more enzymatic activity, whereas lighter fibers indicated little to no enzymatic activity.

Periodic acid-Schiff staining was performed by incubating slides in 0.5% periodic acid for 10 minutes, washing in 3 changes of distilled water, staining in Schiff’s reagent for 5 minutes, and washing in running tap water for 5 minutes to develop color. Slides were then counterstained in hematoxylin for 1 minute to visualize nuclei, and rinsed in running tap water for 1 minute before bluing in 0.1% ammonium water for 1 minute. Slides were rinsed in running tap water for 1 minute before dehydration through 3 changes each of 95% and 100% alcohol, and xylenes before mounting with Cytoseal (Avantor). Glycogenic components (glycogen, glycoproteins, glycolipids) were stained with a magenta hue at the site of accumulation.

Immunofluorescent staining was used to label skeletal muscle fiber subtypes (type 1, 2A, 2B, and 2X) in 10–11-month-old TA samples (n = 5 mice per experimental group). These sections were blocked in PBS + 0.1% Tween-20 + 10% normal goat serum for 1 hour at room temperature prior to primary antibody incubation. Slides were incubated with primary antibodies against laminin (1:500, Sigma, L9393) to stain for basal membranes and for IgG2b (1:200, DSHB, BA-D5-s) for type 1 slow myosin, IgG1 (1:200, DSHB, SC-71-s) for type IIA myosin, and IgM (1:200, DSHB, BF-F3-c) for type IIB myosin overnight at 4°C. Slides were incubated with secondary antibodies against IgG (1:200, Alexa fluor 647), IgG2b (1:200, DyLight 405), IgG1 (1:200, Alexa fluor 568), and IgM (1:200, Alexa fluor 488) for 1 hour at room temperature. Slides were washed with PBS + 0.1% Tween-20 after each blocking and incubation step, and mounted with *Vectashield* mounting medium (without DAPI, VectorLabs). All histological slides were stored in a dark chamber until imaging and semi-automated quantification.

#### Image Acquisition and Morphometric Analyses

Whole muscle sections were imaged by a Keyence BZ-X810 microscope and stitched at 20× magnification for all stains. Stitched images were saved as a 24-bit TIFF without compression for morphometric analysis after image acquisition. H&E-, modified Gömöri Trichrome-, and PAS-stained slides were imaged in brightfield view with an optimized exposure time between 1/200 and 1/250 seconds. Myopathic features (rimmed vacuoles, fiber splitting, internal nuclei, and whorled fibers) in TA muscles from modified Gömöri Trichrome staining were quantified in FIJI. Fibers containing myopathic features were quantified as a proportion of the total fiber count per muscle section, consistent with quantifications from patient biopsies.^1, 2, 14, 23, 32^

The Keyence BZ-X810 microscope was configured with 3 color filters for immunostaining images, (1) DAPI (blue) excitation wavelength (ET) 360/40nm, (2) GFP (green), ET 470/40nm, (3) mCherry, Texas Red, ET 560/40nm. All immunofluorescent images were captured in multichannel view with automated focus and exposure settings used to optimize sharpness and light intensity (with limited photobleaching). The optimized exposure time for immunofluorescent fiber type staining was 0.5 seconds. Immunofluorescent images for fiber type staining were individually captured in three color channels and subsequently merged and stitched from multiple tiles. Image processing for immunofluorescent fiber type staining involved: (1) brightness and contrast enhancement to remove background noise, (2) fiber detection based on laminin fiber outlining, and (3) double extraction of green (type IIB) fibers and red (type IIA) fibers using the Keyence BZ-H4CM macro cell count application. The macro cell count application allowed for a semi-automated quantification of fiber type composition based on set conditions that (1) recognized green versus red color hues, (2) filled gaps between fibers as an exclusion parameter, and (3) allowed for manual removal of false positive artifacts that picked up fluorescent antibody signal. Once these conditions were set and exclusion parameters were modified for each sample, the macro-cell count program generated an analysis of the total cross-sectional area (CSA) for each whole muscle section as well as the proportion of positive green or red areas (μm^2^) relative to the total CSA.

### Transmission Electron Microscopy

Following harvest, a small piece of TA muscle was fixed with 2.5% glutaraldehyde, 2.0% *para*-formaldehyde in 100 mM cacodylate buffer pH 7.2 for 2 h at 4°C, then post-fixed with 1% osmium tetroxide in 1.25% potassium ferrocyanide. Samples were then *en-bloc* stained with 2% uranyl acetate in 0.05 M maleate buffer pH 5.2 overnight at room temperature, followed by serial dehydrations with ethanol, then embedded in resin at 60°C for 48 hours. 60 nm sections were obtained using a diamond knife on a Leica UC67 Ultramicrotome and observed at 120 kV on a Tecnai G2 Spirit BioTWIN Transmission Electron Microscope. Micrographs were captured using a digital camera from Advanced Microscopy Techniques (AMT).

### Statistical Analyses and *p*-value Detection

All statistical analyses were performed using Graph Pad Prism 10.4.1 (GraphPad Software, Boston, MA). The Shapiro-Wilk test was used to assess normality and Levene’s test to assess equal variance across each dataset. Once assumptions of normality and variance were met, unpaired *t*-tests were conducted to assess if there was a statistically significant difference between the mean values of observed properties in *Adss1*^WT/WT^ and *Adss1*^KO/KO^ mice. Given that all experiments compared two independent groups with one metric dependent variable, unpaired *t*-tests were the most suitable method for analysis. For longitudinal studies, multiple unpaired *t*-tests were used to detect *p*-value significance at each timepoint. *p-*values of 0.05 were considered statistically significant, and levels of significance were denoted as \**p* ≤ 0.05, \*\**p* ≤ 0.01, and \*\*\**p* ≤ 0.001.

## ACKNOWLEDGEMENTS

The authors would like to thank and acknowledge the *ADSS1* family community for their participation, support and inspiration for research on *ADSS1* myopathy, and especially Priyanka Kakkar and Naveen Baweja for their pivotal role in spearheading this research and for their tireless efforts in assembling a dedicated international team of researchers and clinicians. We are grateful to Prof. Emma Rybalka at Victoria University for her valuable discussions and scientific insights throughout the project. Profound thanks to Prof. Ling Li at City of Hope Hospital for providing the floxed *Adss1* mice used in these studies, which were instrumental in enabling this work. We also thank Dr. Michael Lawlor for his generous and expert advice on pathology and Dr. Matthias Lambert for kindly sharing his protocol and expertise in fiber typing. Electron microscopy was performed utilizing the resources and assistance of the HMS Electron Microscopy Core Facility and Sanger sequencing was performed utilizing services of the Boston Children’s Hospital Molecular Genetics Core Facility.

## COMPETING INTERESTS

A.H.B. reports consulting income from Astellas Pharma and has equity in Kinea Bio. The remaining authors have no competing interests to declare.

## FUNDING

This work was funded in part by generous philanthropic support from the Lee and Penny Anderson Family Foundation, the Muscular Dystrophy Association (USA), and generous in-kind support for mouse acquisition and derivation from the Cure ADSSL1 Foundation, and Rich Horgan and Cure Rare Disease. Support for molecular genetic analyses was provided by the Boston Children’s Hospital Intellectual and Developmental Disabilities Research Center Molecular Genetics Core Facility supported by P50HD105351 from the Eunice Kennedy Shriver National Institute of Child Health and Human Development of NIH. M.E.K. was supported by the Harvard College Research Program and K.M.Y. was supported by the Boston Children’s Hospital Developmental Neurology Training Grant T32NS007473 from the National Institute of Neurological Disorders and Stroke of NIH.

