## Supplemental Information for "Loss of adenylosuccinate synthetase 1 in mice recapitulates features of *ADSS1* myopathy"

| **Page** | **Content** |
| --- | --- |
| 1 | Table of contents |
| 2 | *Adss2* mRNA expression |
| 3 | EDL and soleus masses |
| 4 | Force frequency curves |
| 5 | *Ex vivo* fatigue analysis for soleus |
| 6 | Myopathic features by histology |
| 7 | Tibialis Anterior fiber composition |
| 8 | Focal brown puncta in NADH-TR stain |

**SUPPORTING FIGURES**

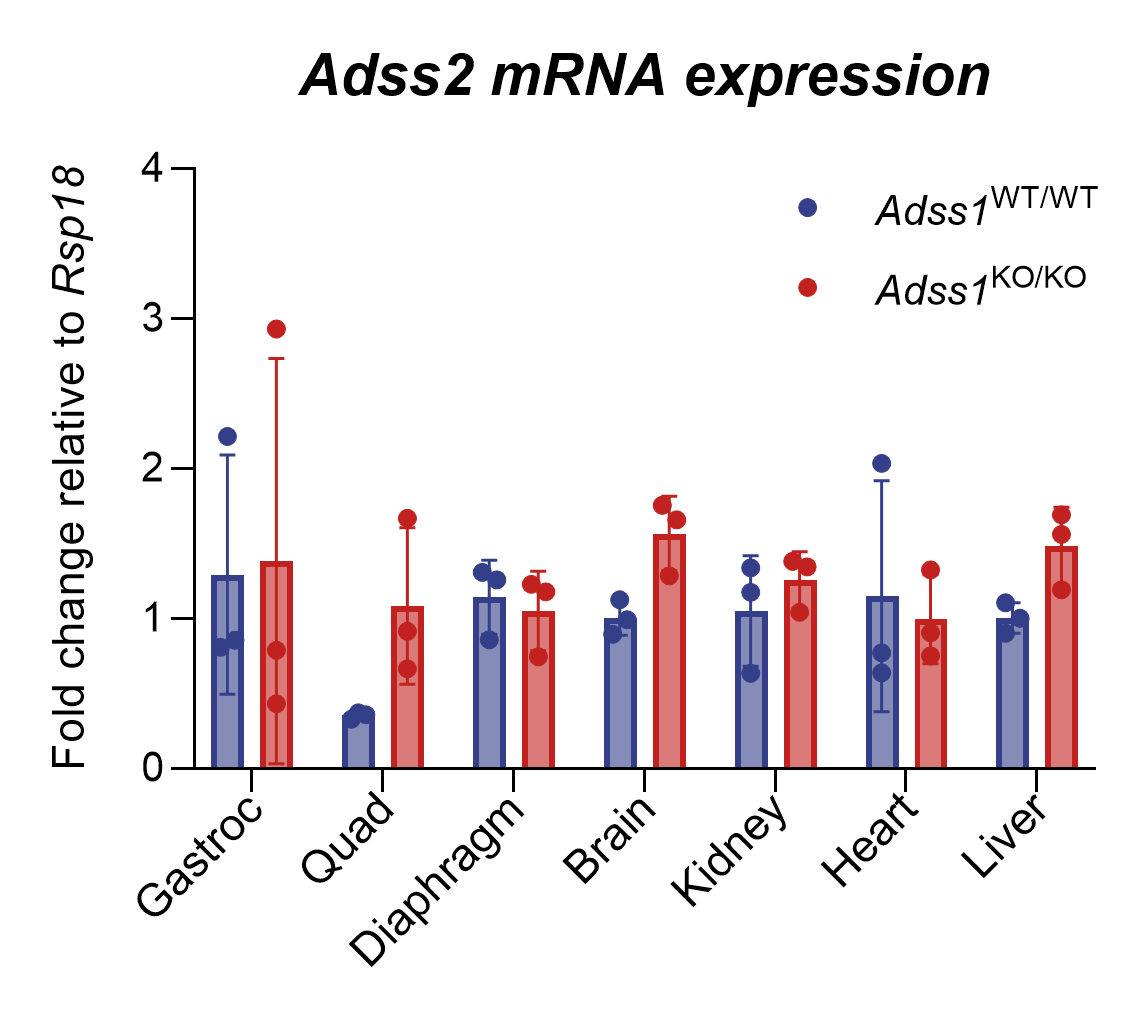
**FIGURE S1**

**Figure S1** mRNA expression levels of *Adss2*, the non-muscle isoform of AdSS. Quantitative PCR indicates that mRNA expression levels are comparable across skeletal muscle tissue and various organs for *Adss1*^KO/KO^ and *Adss1*^WT/WT^ (n = 3 biological replicates). No difference was statistically significant.

**FIGURE S2**

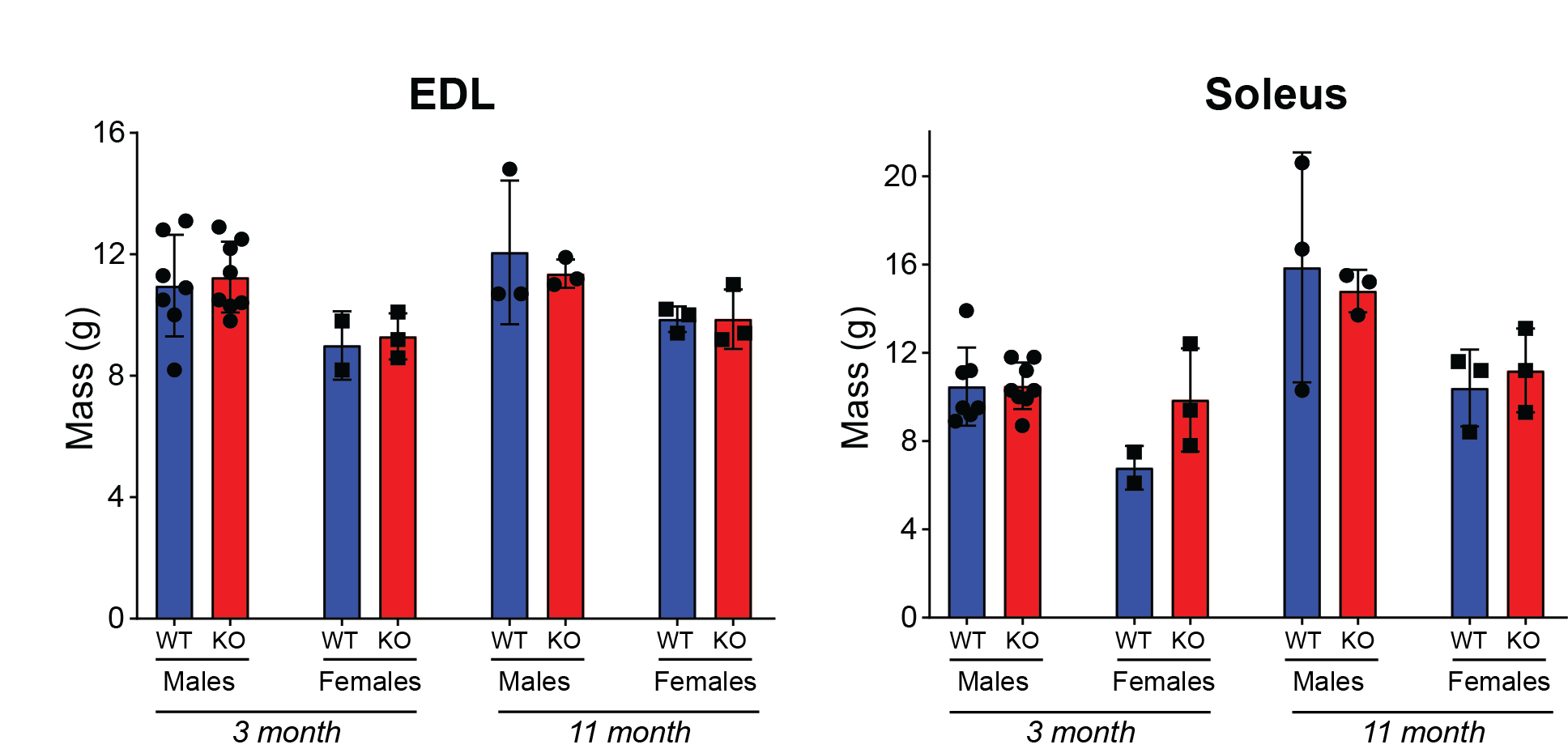

**Figure S2** Mass of EDL and soleus of *Adss1*^KO/KO^ and *Adss1*^WT/WT^ mice at 3 and 11 months of age. No significant difference was observed between the two genotypes.

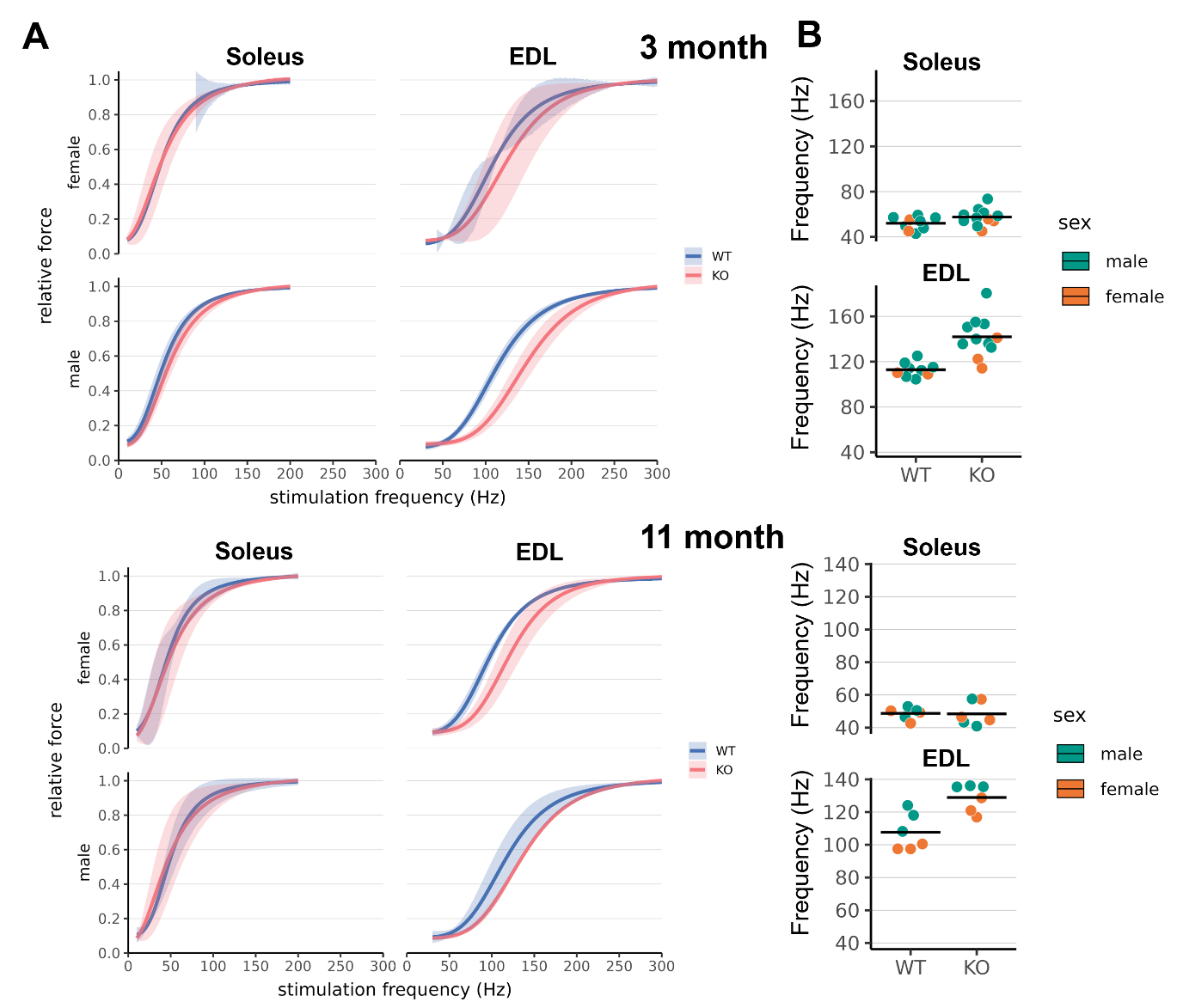
**FIGURE S3**

**Figure S3 (A)** Force-frequency curves of soleus and EDL of *Adss1*^KO/KO^ and *Adss1*^WT/WT^ mice at 3 and 11 months of age. The force-frequency relationship of *Adss1*^KO/KO^ EDL is shifted to right. **(B)** Inflection point frequency of the curves presented in (A). No significant different was observed between the two genotypes.

**FIGURE S4**

**Fatigue Analysis for Soleus**

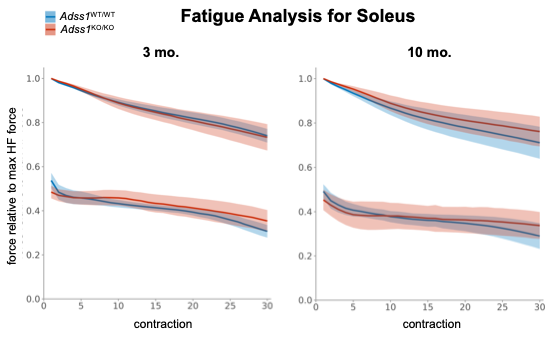

High frequency

High frequency

Low frequency

Low frequency

**Figure S4** *Ex vivo* fatigue analysis on *Adss1*^KO/KO^ and *Adss1*^WT/WT^ soleus muscles measuring the force-frequency relationship between contractile force generated by each soleus from repeated stimuli, relative to their peak force (**Fig. 3**). No difference in contractile force observed between *Adss1*^KO/KO^ and *Adss1*^WT/WT^ soleus muscles after fatigue at 3 months (n = 8) or 10 months (n = 6). Curves plot the mean force, with ribbons representing the 95% confidence interval.

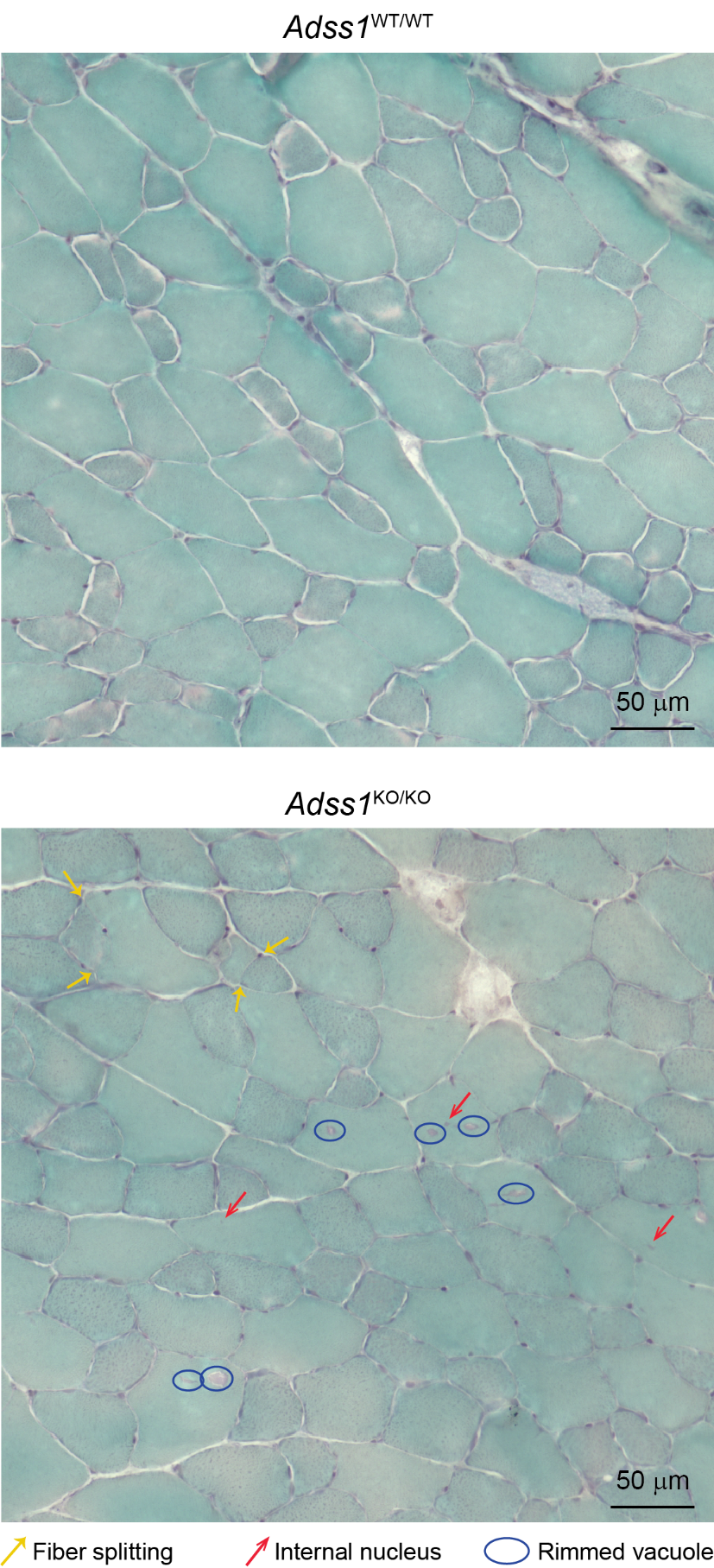
**FIGURE S5**

**Figure S5** *(above)* Representative images of modified Gömöri Trichrome staining of TA sections of 11-month-old *Adss1*^WT/WT^ and *Adss1*^KO/KO^ mice. Many myopathic features were observed in *Adss1*^KO/KO^ sections, including fiber splitting (yellow arrows), internal nuclei (red arrows), and rimmed vacuoles (blue circle). While these features were occasionally seen in wild-type sections, they were much more prevalent in *Adss1*^KO/KO^ sections.

**FIGURE S6**

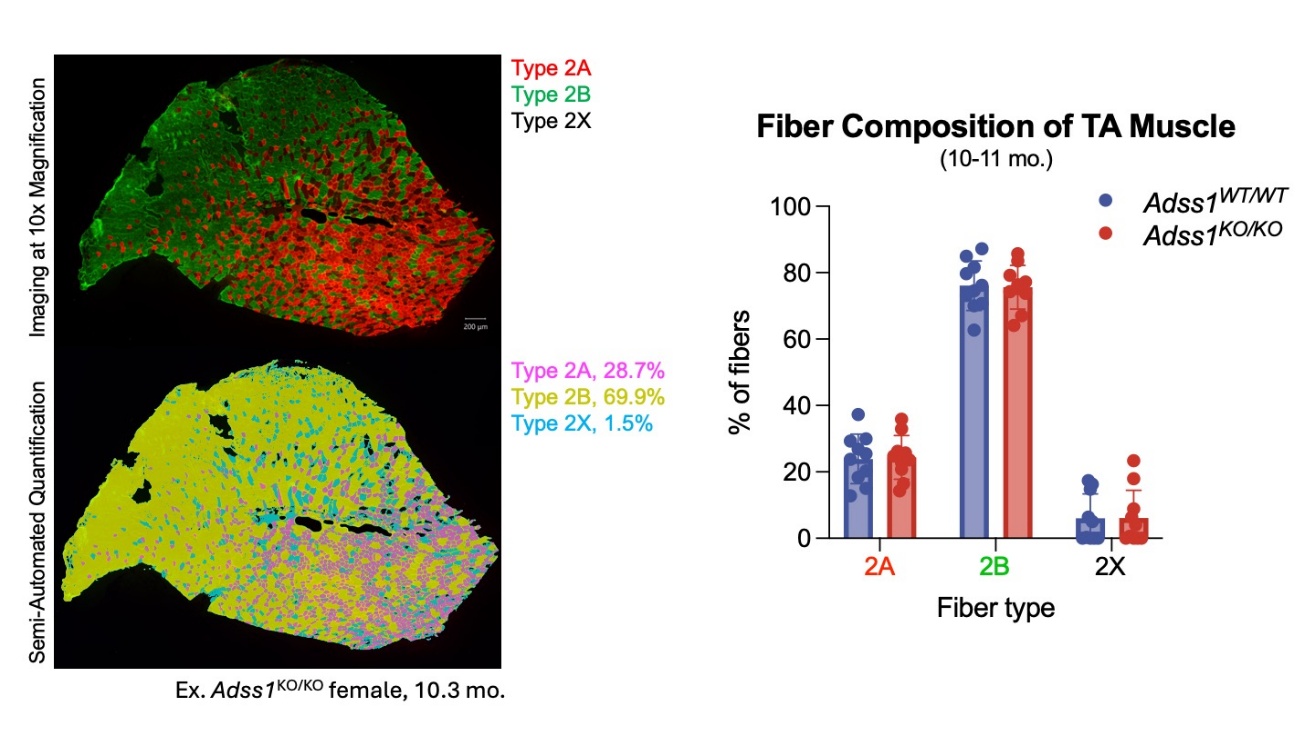

**A**

**B**

**Figure S6 (A)** Semi-automated quantification of fiber typing. Example shown is a representative image of *Adss1*^KO/KO^ female tibialis anterior (TA) muscle, 10.3 months old. **(B)** Fiber type composition of TA muscle (n = 5) measured by the proportion of immunofluorescent areas corresponding to fiber type (%) to total cross-sectional area. (ns, multiple unpaired *t*-tests).

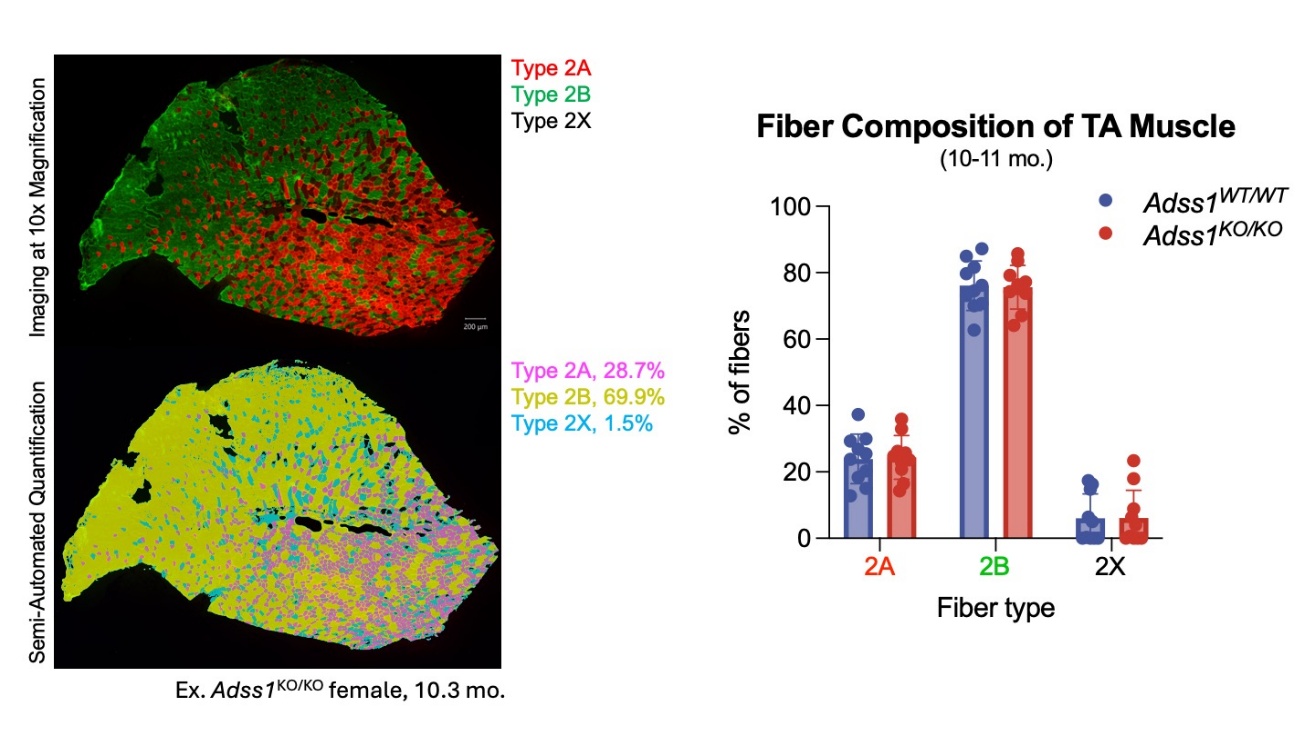

**A**

**B**

**FIGURE S7**

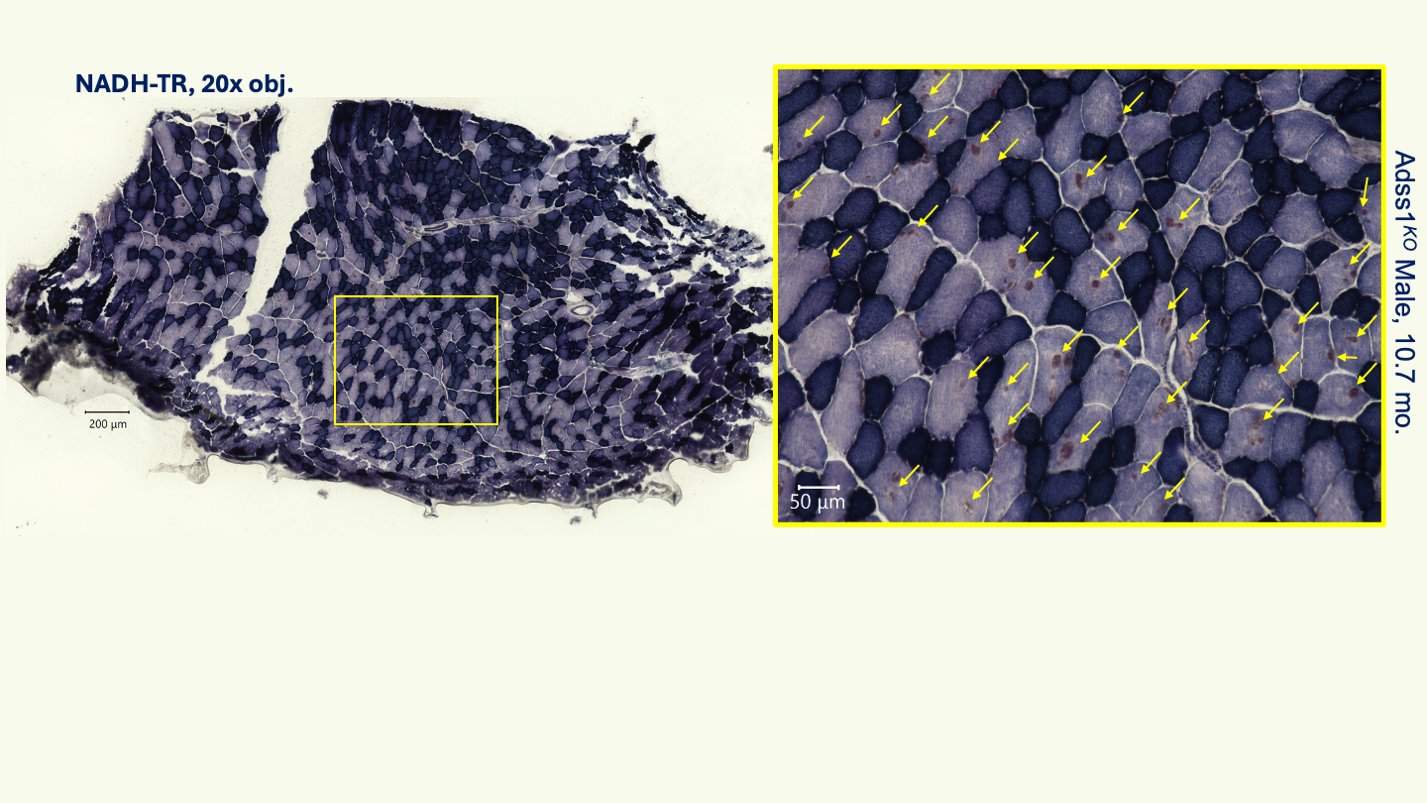

**Figure S7** NADH-TR staining reveals varying levels of oxidative activity associated with fiber type. Oxidative fibers appear as dark purple and glycolytic fibers appear as pale purple. Focal brown deposits are indicated with arrows and appear to co-localize to areas with fiber damage, observed as rimmed vacuoles in modified Gömöri Trichrome staining. Imaged at 20x magnification.
